# Cell cycle stage classification using phase imaging with computational specificity

**DOI:** 10.1101/2021.11.05.467526

**Authors:** Yuchen R. He, Shenghua He, Mikhail E. Kandel, Young Jae Lee, Chenfei Hu, Nahil Sobh, Mark A. Anastasio, Gabriel Popescu

## Abstract

Traditional methods for cell cycle stage classification rely heavily on fluorescence microscopy to monitor nuclear dynamics. These methods inevitably face the typical phototoxicity and photobleaching limitations of fluorescence imaging. Here, we present a cell cycle detection workflow using the principle of phase imaging with computational specificity (PICS). The proposed method uses neural networks to extract cell cycle-dependent features from quantitative phase imaging (QPI) measurements directly. Our results indicate that this approach attains very good accuracy in classifying live cells into G1, S, and G2/M stages, respectively. We also demonstrate that the proposed method can be applied to study single-cell dynamics within the cell cycle as well as cell population distribution across different stages of the cell cycle. We envision that the proposed method can become a nondestructive tool to analyze cell cycle progression in fields ranging from cell biology to biopharma applications.

**Teaser:** We present a non-destructive, high-throughput method for cell cycle detection combining label-free imaging and deep learning.

## Introduction

The cell cycle (*1*) is an orchestrated process that leads to genetic replication and cellular division. This precise, periodic progression is crucial to a variety of processes, such as, cell differentiation, organogenesis, senescence, and disease. Significantly, DNA damage can lead to cell cycle alteration and serious afflictions, including cancer (*2*). Conversely, understanding the cell cycle progression as part of the cellular response to DNA damage has emerged as an active field in cancer biology (*3*).

Morphologically, the cell cycle can be divided into interphase and mitosis. The interphase (*1*) can further be divided into three stages: G1, S, and G2. Since the cells are preparing for DNA synthesis and mitosis during G1 and G2 respectively, these two stages are also referred to as the “gaps” of the cell cycle (*4*). During the S stage, the cells are synthesizing DNA, with the chromosome count increasing from 2N to 4N.

Traditional approaches for distinguishing different stages within the cell cycle rely on fluorescence microscopy (*5*) to monitor the activity of proteins that are involved in DNA replication and repair, e.g., proliferating cell nuclear antigen (PCNA) (*6*). A variety of signal processing techniques, including support vector machine (SVM) (*7*), intensity histogram and intensity surface curvature (*8*), level-set segmentation (*9*), and k-nearest neighbor (*10*) have been applied to fluorescence intensity images to perform classification. In recent years, with the rapid development of parallel-computing capability (*11*) and deep learning algorithms (*12*), convolutional neural networks have also been applied to fluorescence images of single cells for cell cycle tracking (*13, 14*). Since all these methods are based on fluorescence microscopy, they inevitably face the associated limitations, including photobleaching, chemical, and phototoxicity, weak fluorescent signals that require large exposures, as well as nonspecific binding. These constraints limit the applicability of fluorescence imaging to studying live cell cultures over large temporal scales (*15*).

Quantitative phase imaging (QPI) (*16*) is a family of label-free imaging methods that has gained significant interest in recent years due to its applicability to both basic and clinical science (*17*). Since the QPI methods utilize the optical path length as intrinsic contrast, the imaging is non-invasive and, thus, allows for monitoring live samples over several days without concerns of degraded viability (*17*). As the refractive index is linearly proportional to the cell density (*18*), independent of the composition, QPI methods can be used to measure the non-aqueous content (dry mass) of the cellular culture (*19*). In the past two decades, QPI has also been implemented as a label-free tomography approach for measuring 3D cells and tissues (*20-27*). These QPI measurements directly yield biophysical parameters of interest in studying neuronal activity (*28*), quantifying sub-cellular contents (*29*), as well as monitoring cell growth along the cell cycle (*30-32*). Recently, with the parallel advancement in deep learning, convolutional neural networks were applied to QPI data as universal function approximators (*33*) for various applications (*34*). It has been shown that deep learning can help computationally substitute chemical stains for cells (*35*) and tissues (*36*), extract biomarkers of interest (*37*), enhance imaging quality (*38*), as well as solve inverse problems (*39*).

In this article, we present a new methodology for cell cycle detection that utilizes the principle of phase imaging with computational specificity (PICS) (*37*). Our approach combines spatial light interference microscopy (SLIM) (*40*), a highly sensitive QPI method, with recently developed deep learning network architecture E-U-Net (*41*). We demonstrate on live cell cultures that the proposed method classifies cell cycle stages solely using SLIM images as input. The signals from the fluorescent ubiquitination-based cell cycle indicator (FUCCI) (*42*) were only used to generate ground truth annotations during the deep learning training stage. Unlike previous methods that perform single-cell classification based on bright-field and dark-field images from flow cytometry (*43*) or phase images from ptychography (*44*), our method can classify all adherent cells in the field of view and perform longitudinal studies over many cell cycles. Evaluated on a test set consisting of 408 unseen SLIM images (over 10,000 cells), our method achieves F-1 scores over 0.75 for both the G1 and S stage. For the G2/M phase, we obtained a lower score of 0.6, likely due to the round cells going out of focus in the M-stage. Using the classification data outputted by our method, we created binary maps that were used back into the QPI (input) images to measure single cell area, dry mass, and dry mass density for large cell populations in the three cell cycle stages. Because our SLIM imaging is nondestructive, all individual cells can be monitored over many cell cycles without loss of viability. We envision that our proposed method can be extended to different cell lines and other QPI imaging modalities for high throughput and nondestructive cell cycle analysis, thus, eliminating the need for cell synchronization.

## Results

The experiment setup is illustrated in Fig. 1. We utilized spatial light interference microscopy (SLIM) (*40*) to acquire the quantitative phase map of live HeLa cells prepared in six-well plates. By adding a QPI module to an existing phase contrast microscope, SLIM modulates the phase delay between the incident field and the scattered field, and an optical pathlength map is then extracted from four intensity images via phase-shifting interferometry (*16*). Due to the common-path design of the optical system, we were able to acquire both the SLIM signals and epi-fluorescence signals of the same field of view (FOV) using a shared camera. Figure 1B shows the quantitative phase map of live HeLa cell cultures using SLIM.

**Fig. 1.**
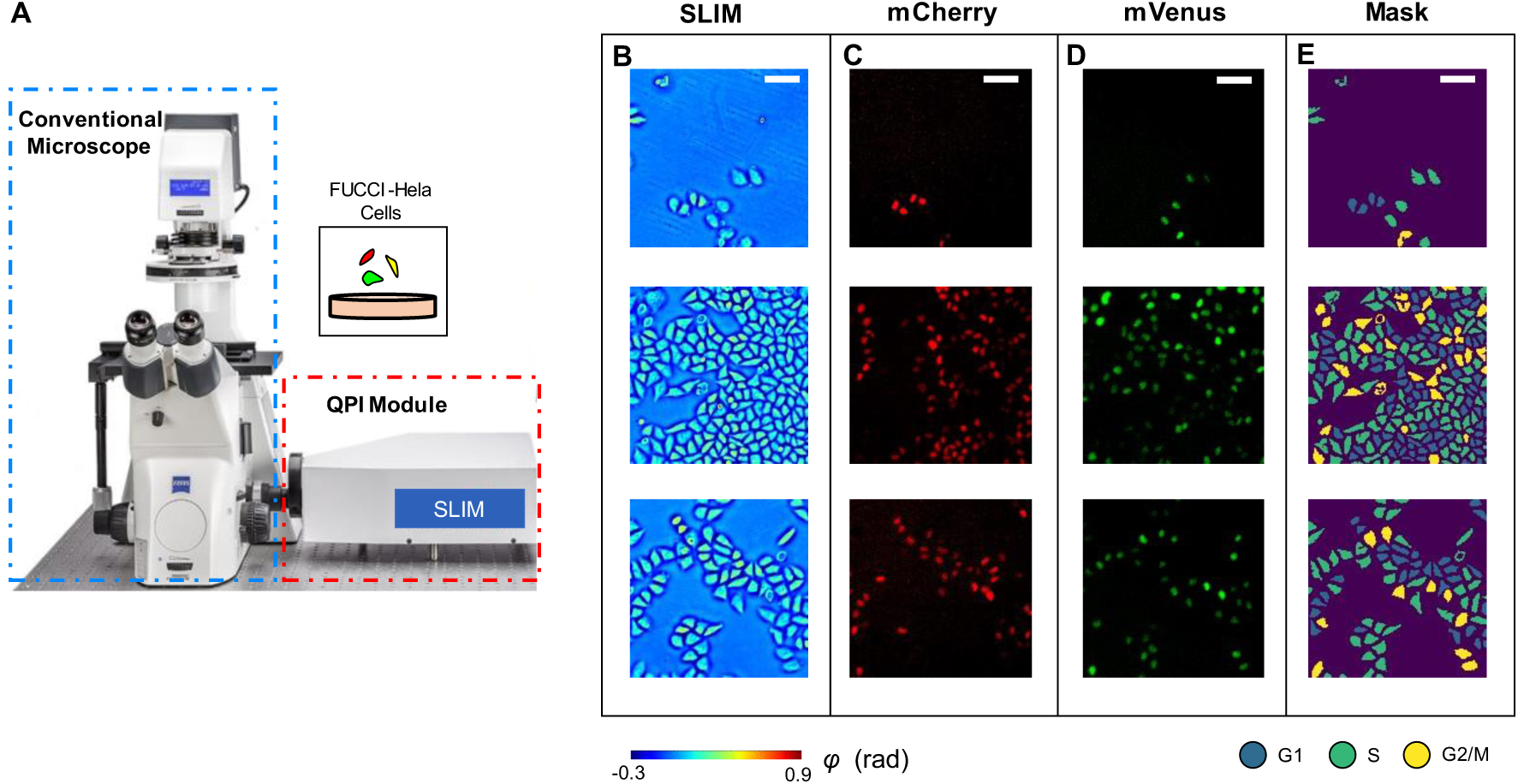
Schematic of the imaging system. (**A**) The SLIM module was connected to the side port of an existing phase contrast microscope. This setup allows us to take co-localized SLIM images and fluorescence images by switching between transmission and reflection illumination. (**B**) Measurements of HeLa cells. (**C**) mCherry fluorescence signals. d. mVenus fluorescence signals. e. Cell cycle phase masks generated by using adaptive thresholding to combine information from all three channels. Scale bar is 100 μm.

To obtain an accurate classification between the three stages within one cell cycle interphase (G1, S, and G2), we used HeLa cells that were encoded with fluorescent ubiquitination-based cell cycle indicator (FUCCI) (*42*). FUCCI employs mCherry, an hCdt1-based probe, and mVenus, an hGem-based probe, to monitor proteins associated with the interphase. FUCCI transfected cells produce a sharp triple color-distinct separation of G1, S, and G2/M. Figure 1C and 1D demonstrate the acquired mCherry signal and mVenus signal, respectively. We combined the information from all three channels via adaptive thresholding to generate a cell cycle stage mask (Fig. 1E). The procedure of sample preparation and mask generation is presented in detail in the Methods section and Fig. S1.

### Deep Learning

With the SLIM images as input and the FUCCI cell masks as ground truth, we formulated the cell cycle detection problem as a semantic segmentation task and trained a deep neural network to infer each pixel’s category as one of the “G1”, “S”, “G2/M”, or background labels. Inspired by the high accuracy reported in previous works (*41*), we used the E-U-Net (Fig. 2A) as our network architecture. The E-U-Net architecture upgraded the classic U-Net (*45*) by swapping its original encoder layers with a pre-trained EfficientNet (*46*). Since the EfficientNet was already trained on the massive ImageNet dataset, it provided more sophisticated initial weights than the randomly initialized layers from the scratch U-Net as in previous approaches (*37*). This transfer learning strategy enables the model to utilize “knowledge” of feature extraction learned from the ImageNet dataset, achieving faster convergence and better performance (*41*). Since EfficientNet was designed using a compound scaling coefficient, it is still relatively small in size. Our trained network used EfficientNet-B4 as the encoder and contained 25 million trainable parameters in total.

**Fig. 2.**
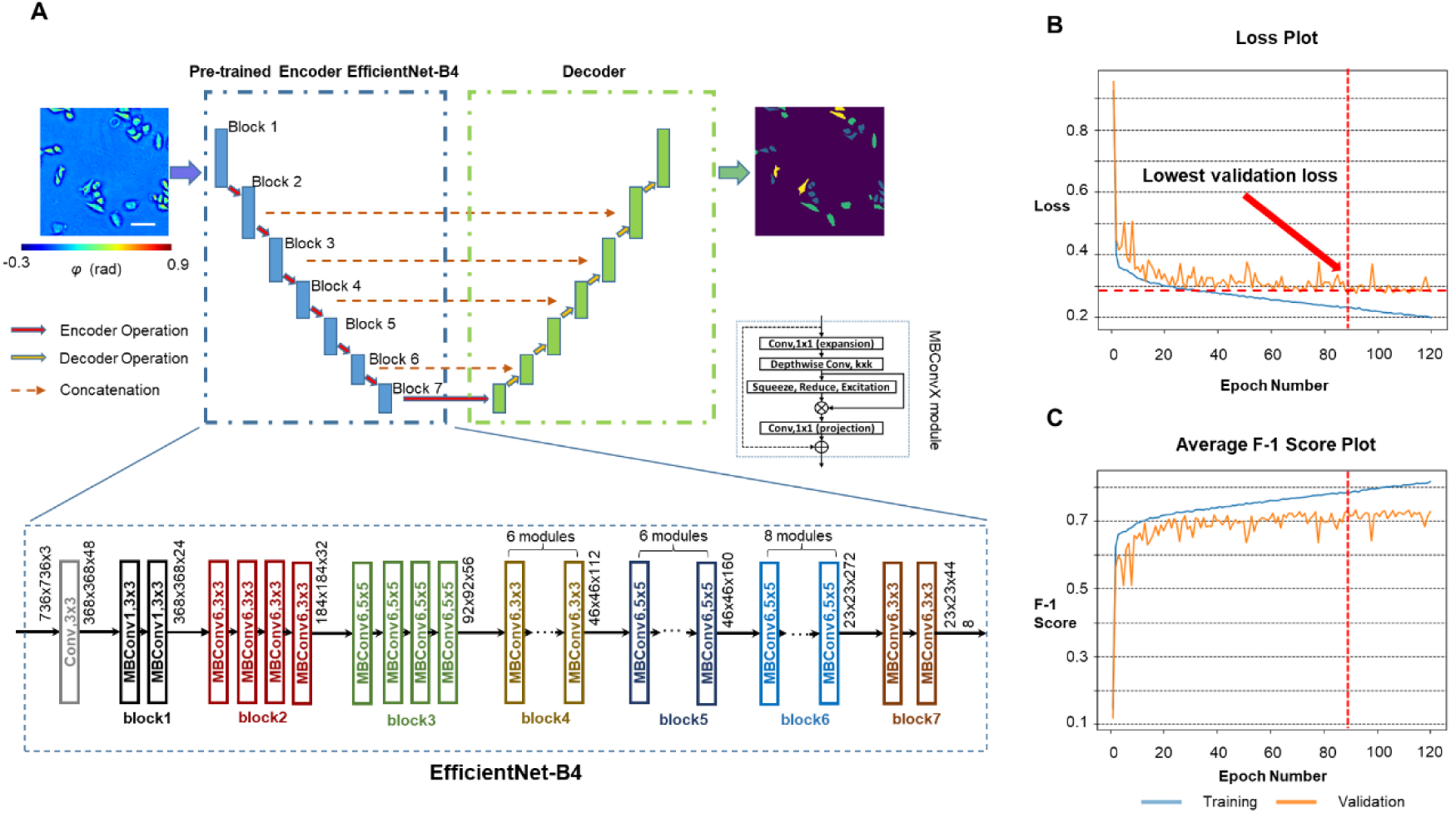
PICS training procedure. (**A**) We used a network architecture called the E-U-Net that replaces the encoder part of a standard U-Net with the pre-trained EfficientNet-B4. Within the encoder path, the input images were downsampled 5 times through 7 blocks of encoder operations. Each encoder operation consists of multiple MBConvX modules that consist of convolutional layers, squeeze and excitation, and residual connections. The decoder path consists of concatenation, convolution and upsampling operations. (**B**) The model loss values on the training dataset and the validation dataset after each epoch. We picked the model checkpoint with the lowest validation loss as our final model and used it for all analysis. (**C**) The model’s average F-1 score on the training dataset and the validation dataset after each epoch.

We trained our E-U-Net with 2,046 pairs of SLIM images and ground truth masks for 120 epochs. The network was optimized by an Adam optimizer (*47*) against the sum of the DICE loss (*48*) and the categorical focal loss (*49*). After each epoch, we computed the model’s loss on both the training set and the validation set, which consists of 408 different image pairs (Fig. 2B). The weights of parameters that make the model achieve the lowest validation loss were selected and used for all verification and analysis. The training procedure is described in Methods.

### PICS Performance

After training the model, we evaluated its performance on 408 unseen SLIM images from the test dataset. The test dataset was selected from wells that are different from the ones used for network training and validation during the experiment. Figure 3A shows randomly selected images from the test dataset. Figure 3B and 3C show the corresponding ground truth cell cycle masks and the PICS cell cycle masks, respectively. It can be seen that the trained model was able to identify the cell body accurately.

**Fig. 3.**
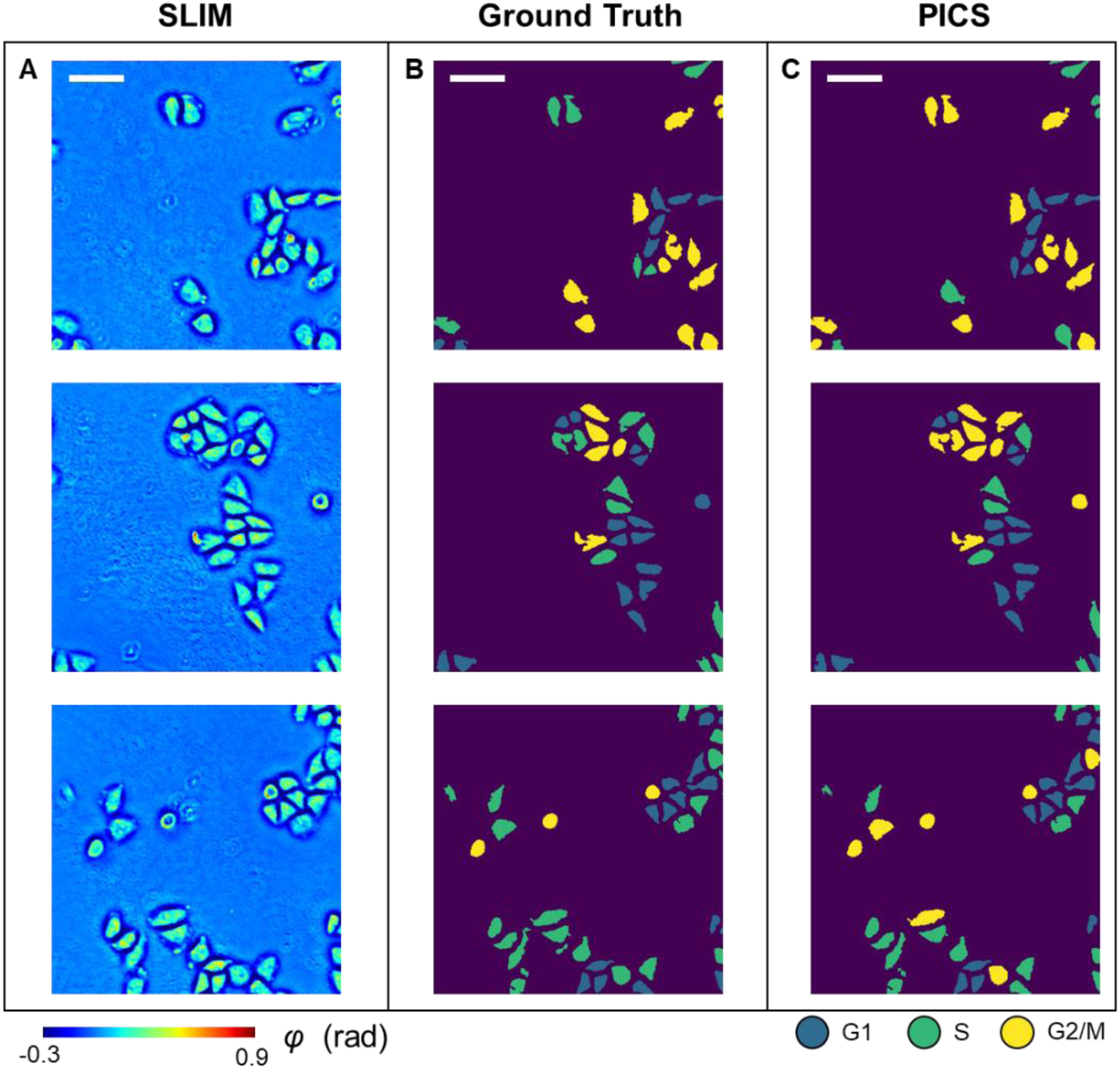
PICS results on the test dataset. (**A**) SLIM images of Hela cells from the test dataset. (**B**) Ground truth cell cycle phase masks. (**C**) PICS-generated cell cycle phase masks. Scale bar is 100 μm.

We reported the raw performance of our PICS methods in Fig. S2, with pixel-wise precision, recall, and F1-score for each class. However, we noticed that these metrics did not reflect the performance in terms of the number of cells. Thus, we performed a post-processing step on the inferred masks to enforce particle-wise consistency, as detailed in Methods and Fig. S3. After this post-processing step, we evaluated the model’s performance on the cellular level and produced the cell count-based results shown in Fig. 4. Figure 4A shows the histogram of cell body area for cells in different stages, derived from both the ground truth masks and the prediction masks. Figures 4B and 4C show similar histograms of cellular dry mass and dry mass density, respectively. The histograms indicated that there is a close overlap between the quantities derived from the ground truth masks and the prediction masks. The cell-wise precision, recall, and F-1 score for all three stages are shown in Fig. 4D. Each entry is normalized with respect to the ground truth number of cells in that stage. Our deep learning model achieved over 0.75 F-1 scores for both the G1 stage and the S stage, and a 0.6 F-1 score for the G2/M stage. The lower performance for the G2/M stage is likely due to the round cells going out of focus during mitosis.

**Fig. 4.**
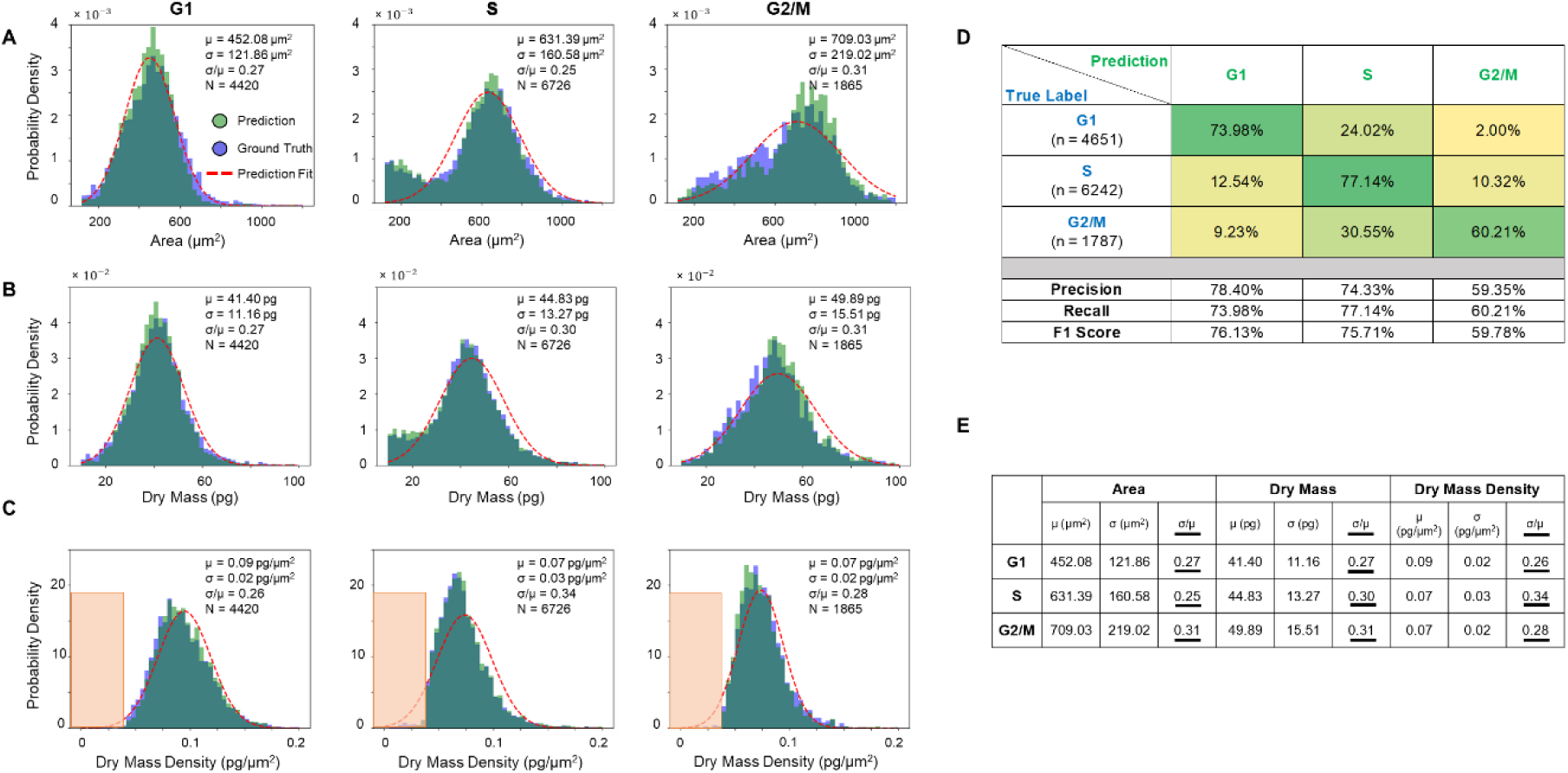
PICS performance on the test dataset. (**A-C**) Histograms of cell area, dry mass and dry mass density for cells in G1, S, and G2/M, generated by the ground truth mask (in blue) and by PICS (in green). A Gaussian distribution (in red) was fitted to the PICS results. (**D**) Confusion matrix for PICS inference on the test dataset. (**E**) Mean, standard deviation and their ratio (underlined for visibility) of cell area, dry mass and dry mass density obtained from the fitted Gaussian distribution.

We calculated the means and standard deviations of the best fit Gaussian for the area, dry mass, and dry mass density distributions for populations of cells in each of the three stages: G1 (N=4,430 cells), S (N=6,726 cells), and G2/M (1,865 cells). The standard deviation divided by the mean, *σ*/*μ*, is a measure of the distribution spread. These values are indicated in each panel of Figs. 4A-C and summarized in Fig. 4E. We note that the G1 phase is associated with distributions that are most similar to a Gaussian. It is interesting that the S-phase exhibits a bimodal distribution in both area and dry mass, indicating the presence of a subpopulation of smaller cells at the end of G1 phase. However, the dry mass density even for this bimodal population becomes monomodal, suggesting that the dry mass density is a more uniformly distributed parameter, independent of cell size and weight. Similarly, the G2/M area and dry mass distributions are skewed toward the origin, while the dry mass density appears to have a minimum value of ∼0.0375 pg/μm^2^ (within the orange rectangles). Interestingly, early studies of fibroblast spreading also found that there is a minimum value for the dry mass density that cells seem to exhibit (*50*).

### PICS Application

The PICS method can be applied to track the cell cycle transition of single cells, nondestructively. Figure 5A shows the time-lapse SLIM measurements and PICS inference of HeLa cells. The time increment was roughly two hours between two measurements and the images at *t* = 2, 6, 10, and 14 hours were displayed in Fig. 5A. Our deep learning model has not seen any of these SLIM images during training. The comparison between the SLIM images and the PICS inference showed that the deep learning model produced accurate cell body masks and assigned viable cell cycle stages. We showed in Fig. 5B-C the results of manually tracking two cells in this field of view across 16 hours and using the PICS cell cycle masks to compute their cellular area and dry mass. Figure 5B demonstrates the cellular area and dry mass change for the cell marked by the red rectangle. We observed an abrupt drop in both the area and dry mass around *t* = 8 hours, at which point the mother cell divides into two daughter cells. The PICS cell cycle mask also captured this mitosis event as it progressed from the “G2/M” label to the “G1” label. We observed a similar drop in Fig. 5C after 14 hours due to mitosis marked by the orange rectangle. Figure 5C also shows that the cell continues growing before *t* = 14 hours and the PICS cell cycle mask progressed from the “S” label to the “G2/M” label correspondingly. Note that this long-term imaging is only possible due to the nondestructive imaging allowed by SLIM.

**Fig. 5.**
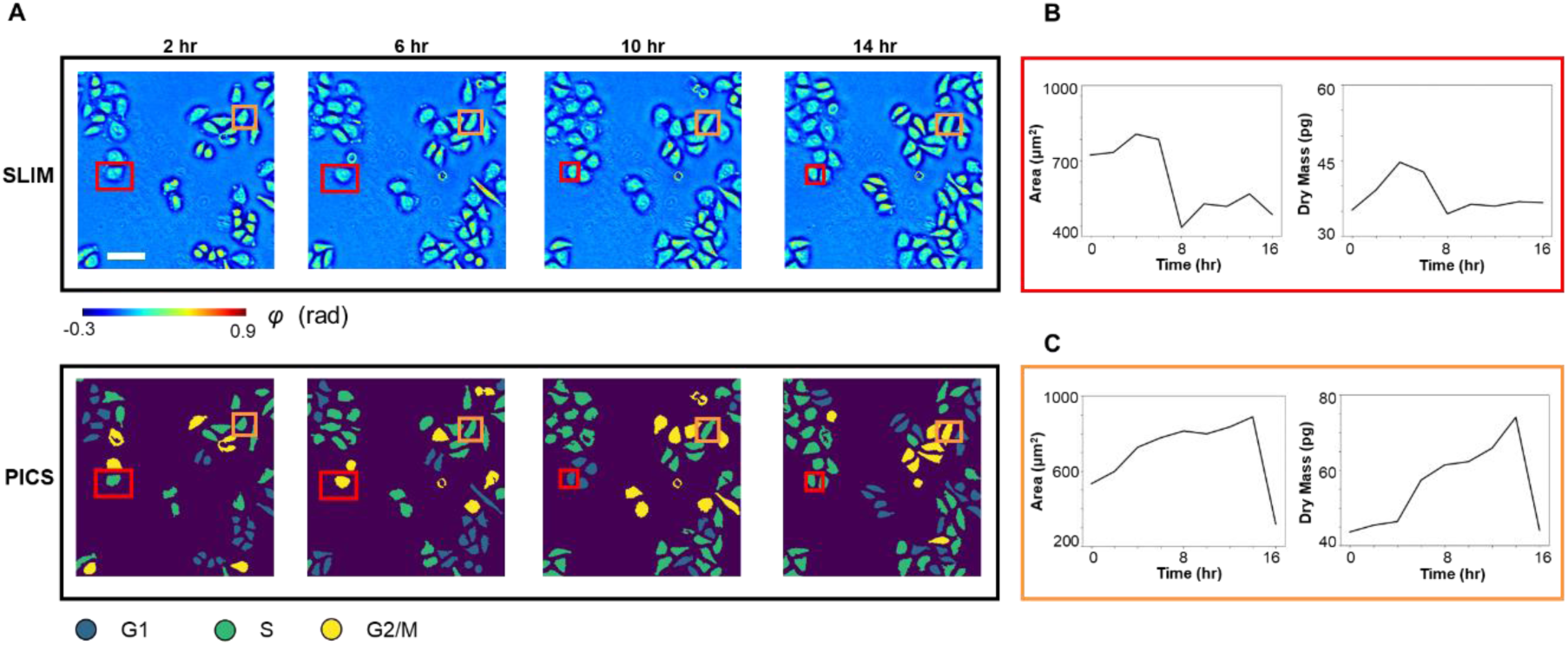
PICS on time lapse of FUCCI-Hela cells. (**A**) SLIM images and PICS inference of cells measured at 2, 6, 10, and 14 hours. The time interval between imaging is roughly 2 hours. We manually tracked two cells (marked in red and orange). (**B**) Cell area and dry mass change of the cell in the red rectangle, across 16 hours. These values were obtained via PICS inferred masks. We can observe an abrupt drop in cell dry mass and area as the cell divides after around 8 hours. (**C**) Cell area and dry mass change of the cell in orange rectangle, across 16 hours. We can observe that the cell continues growing in the first 14 hours as it goes through G1, S, and G2 phase. It divides between hour 14 and hour 16, with an abrupt drop in its dry mass and cell area. Scale bar is 100 μm.

To further demonstrate that PICS can be used to study the statistical distribution of cells across different stages. The PICS inferred cell area distribution across G1, S, and G2/M is plotted in Fig. 6A, whereby a clear shift between cellular area in these stages can be observed. We performed Welch’s t-test on these three groups of data points. To avoid the impact on p-value due to the large sample size, we randomly sampled 20% of all data points from each group and performed the t-test on these subsets instead. After sampling, we have 884 cells in G1, 1345 cells in S, and 373 cells in G2/M. The p-values are less than 10^−3^, indicating statistical significance. The same analysis was performed on the cell dry mass and cell dry mass density, as shown in Figs. 6B-C. We observed a clear distinction between cell dry mass in S and G2/M as well as between cell dry mass density in G1 and S. These results agree with the general expectation that cells are metabolically active and grow during G1 and G2. During S, the cells remain metabolically inactive and replicate their DNA. Since the DNA dry mass only accounts for a very small factor of the total cell dry mass (*32*), the distinction between G1 cell dry mass and S cell dry mass is less obvious than the distinction between S cell dry mass and G2/M cell dry mass. We also noted that our observation on the cell dry mass density distribution agrees with previous findings (*31*).

**Fig. 6.**
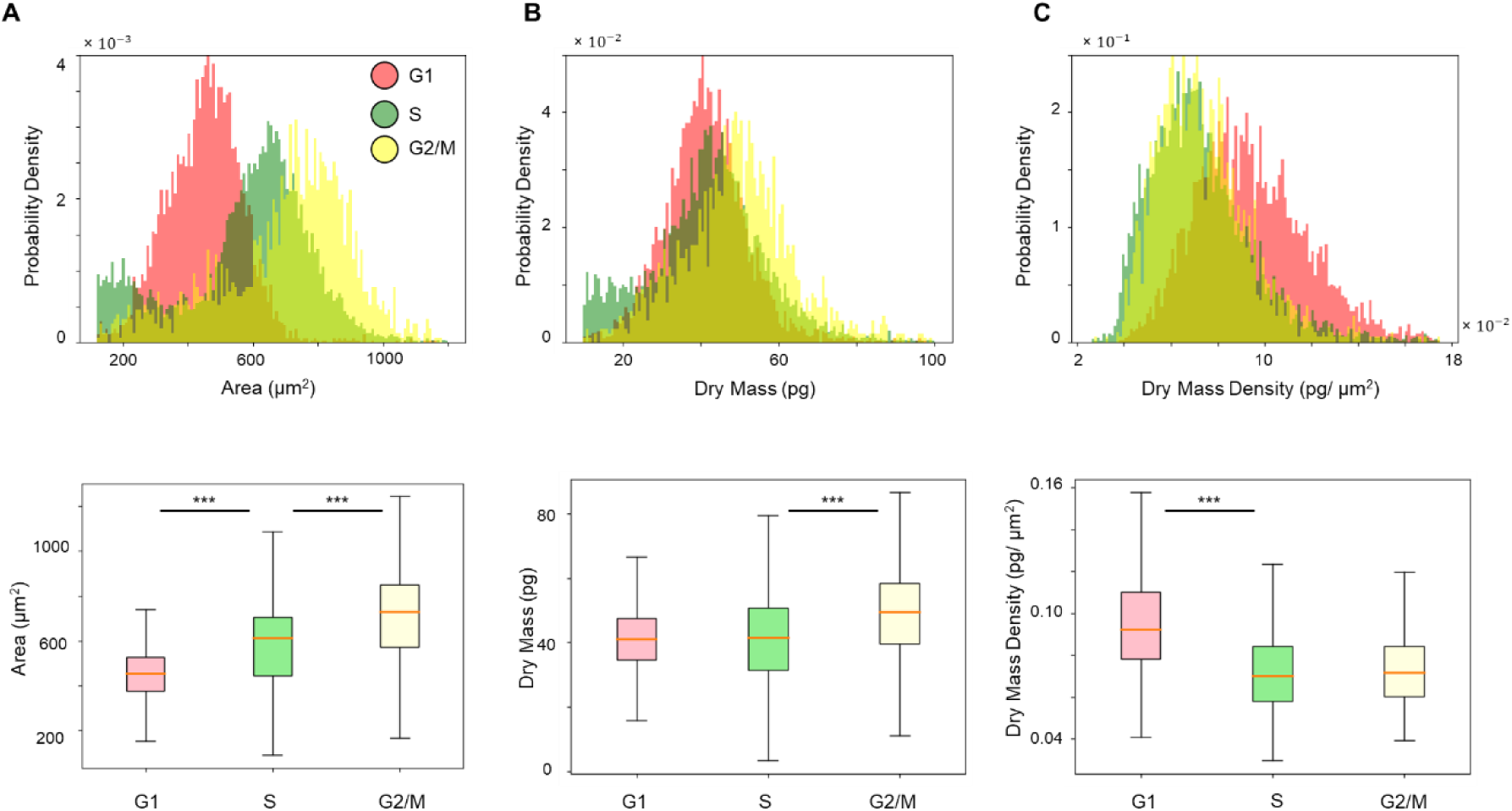
Statistical analysis from PICS inference on the test dataset. (**A**) Histogram and box plot of cell area. The p-value returned from Welch’s t-test indicated statistical significance. (**B**) Histogram and box plot of cell dry mass. The p-value returned from Welch’s t-test indicated statistical significance. (**C**) Histogram and box plot of cell dry mass density. The p-value returned from Welch’s t-test indicated statistical significance comparing cells in G1 and S. The box plot and Welch’s t-test are computed on 20% of all data points in G1, S, and G2/M, randomly sampled. The sample size is 884 for G1, 1345 for S, and 373 for G2/M. Outliers are omitted from the box plot. (*** p < 0.001).

## Discussion

We proposed a PICS-based cell cycle stage classification workflow for fast, label-free cell cycle analysis on adherent cell cultures. Our new method utilizes trained deep neural networks to infer an accurate cell cycle mask from a single SLIM image. The method can be applied to study single-cell growth within the cell cycle as well as compare the cellular parameter distributions between cells in different cell cycle phases.

Compared to many existing methods of cell cycle detection (*7-10, 13, 14, 43, 44, 51*), we believe that our method has three main advantages. First, our method uses a SLIM module, which can be installed as an add-on component to a conventional phase contrast microscope. The imaging cost and setup complexity are much lower compared to previous methods using flow cytometry (*43*), while the user experience remains the same as using a commercial microscope. Significantly, due to the seamless integration with the fluorescence channel on the same field of view, the instrument can collect the ground truth data very easily, while the annotation is automatically performed via thresholding, rather than manually. Second, our method does not rely on fluorescence signals as input. On the contrary, our method is built upon the capability of neural networks to extract label-free cell cycle markers from the quantitative phase map. Thus, the method can be applied to live cell samples over long periods of time without concerns of photobleaching or degraded cell viability due to chemical or phototoxicity. Third, our approach can be applied to large sample sizes consisting of entire fields of views and hundreds of cells. Since we formulated the task as semantic segmentation and trained our model on a dataset containing images with various cell counts, our method worked with FOVs containing up to hundreds of cells. Also, since the U-Net (*45*) style neural network is fully convolutional, our trained model can be applied to images with arbitrary size. Consequently, the method can directly extend to other cell datasets or experiments with different cell confluences, as long as the magnification and numerical aperture stay the same. Since the input imaging data is nondestructive, we can image large cell populations over many cell cycles and study cell cycle phase-specific parameters at the single cell scale. As an illustration of this capability, we measured distributions of cell area, dry mass and dry mass density for populations of thousands of cells in various stages of the cell cycle. We found that the dry mass density distribution drops abruptly under a certain value for all cells, which indicates that live cells require a minimum dry mass density.

During the development of our method, we followed standard protocols in the community(*52*), such as preparing a diverse enough training dataset, properly splitting the training, validation and test dataset, and closely monitoring the model loss convergence to ensure that our model can generalize. We believe PICS-based instruments are well-suited for extending our generalizable method to different cell lines and imaging conditions as the effort to perform extra training is minimal (*37*). Our typical training takes approximately 20 hours, while the inference is performed within 65 ms per frame. Thus, we envision that our proposed workflow is a valuable alternative to the existing methods for cell cycle stage classification and eliminates the need for cell synchronization.

## Materials and Methods

### FUCCI cell and HeLa cell preparation

HeLa/FUCCI(CA)2 (*42*) cells were acquired from RIKEN cell bank and kept frozen in liquid nitrogen tank. Prior to the experiments, we thawed and cultured cells into T75 flasks in Dulbecco’s Modified Eagle Medium (DMEM with low glucose) containing 10% fetal bovine serum (FBS) and incubated in 37°C with 5% CO_2_. When the cells reached 70% confluency, the flask was washed with phosphate-buffered saline (PBS) and trypsinized with 4 mL of 0.25% (w/v) Trypsin EDTA for four minutes. When the cells started to detach, they were suspended in 4 mL of DMEM and passaged onto a glass-bottom six-well plate. HeLa cells were then imaged after two days of growth.

### SLIM imaging

The SLIM system architecture is shown in Fig. 1A. We attached a SLIM module (CellVista SLIM Pro; Phi Optics) to the output port of a phase contrast microscope. Inside the SLIM module, the spatial light modulator matched to the back focal plane of the objective controlled the phase delay between the incident field and the reference field. We recorded four intensity images at phase shifts of 0, *π*/2, π, and 3*π*/2 and reconstructed the quantitative phase map of the sample. We measured both the SLIM signal and the fluorescence signal with a 10×/0.3NA objective. The camera we used was Andor Zyla with a pixel size of 6.5 μm. The exposure time for SLIM channel and fluorescence channel was set to 25 ms and 500 ms, respectively.

### Cellular dry mass computation

We recovered the dry mass as

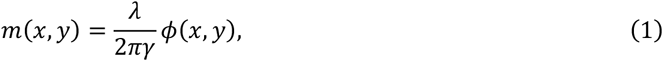

using the same procedure outlined in previous works (*18, 19*). *λ* = 550 *nm* is the central wavelength; *γ* = 0.2 *ml*/*g* is the specific refraction increment, corresponding to the average of reported values (*18, 53*); and *ϕ*(*x, y*) is the measured phase. Equation 1 provides the dry mass density at each pixel, and we integrated over the region of interest to get the cellular dry mass.

### Ground truth cell cycle mask generation

To prepare the ground truth cell cycle masks for training the deep learning models, we combined information from the SLIM channel and the fluorescence channels (Fig. S1A) by applying adaptive thresholding (Fig. S1B). All the code was implemented in Python, using the scikit-image library. We first applied the adaptive thresholding algorithm on the SLIM images to generate accurate cell body masks. Then we applied the algorithm on the mCherry fluorescence images and mVenus fluorescence images to get the nuclei masks that indicate the presence of the fluorescence signals. We took the intersection between the three sets of masks. Following the FUCCI color readout detailed in (*42*), a presence of mCherry signal alone indicates the cell is in G1 stage and a presence of mVenus signal alone indicates the cell is in S stage. The overlapping of both signals indicates the cell is in G2 or M stage. Since the cell mask is always larger than the nuclei mask, we filled in the entire cell area with the corresponding label. We handled the case of no fluorescence signal by automatically labeling them as S because both fluorescence channels yield low-intensity signals only at the start of the S phase (*42*). Before using the mask for analysis, we also performed traditional computer vision operations, e.g., hole filling. on the generated masks to ensure the accuracy of computed dry mass and cell area (Fig. S1C).

### Deep learning model development

We used the E-U-Net architecture (*41*) to develop the deep learning model that can assign a cell cycle phase label to each pixel. The E-U-Net upgraded the classic U-Net (*45*) architecture by swapping its encoder component with a pre-trained EfficientNet (*46*). Compared to previously reported transfer-learning strategies, e.g. utilizing a pre-trained ResNet(*54*) for the encoder part, we believe the E-U-Net architecture is superior since the pre-trained EfficientNet attains higher performance on the benchmark dataset while remaining compact due to the compound scaling strategy (*46*).

The EfficientNet backbone we ended up using for this project was EfficientNet-B4 (Fig. 2A). The entire E-U-Net-B4 model contains around 25 million trainable parameters, which is smaller compared to the number of parameters from the stock U-Net (*45*) and other variations (*55*). We trained the network with 2046 image pairs in the training dataset and 408 image pairs in the validation dataset. Each image contains 736 × 736 pixels. The model was optimized using an Adam optimizer (*47*) with default parameters against the sum of the DICE loss (*48*) and the categorical focal loss (*49*). The DICE loss was designed to maximize the dice coefficient *D* (Equation 2) between the ground truth label (*g*_*i*_) and prediction label (*p*_*i*_) at each pixel. It has been shown in previous works that DICE loss can help tackle class imbalance in the dataset (*56*). Besides DICE loss, we also utilized the categorical focal loss *FL*(*p*_*t*_) (Equation 3). The categorical focal loss extended the cross entropy loss by adding a modulating factor (1 − *p*_*t*_)^*γ*^. It helped the model to focus more on wrong inferences by preventing easily classified pixels dominating the gradient. We tuned the ratio between these two loss values and launched multiple training sessions. In the end we found the model trained against an equally weighted DICE loss and categorical focal loss gave the best results.

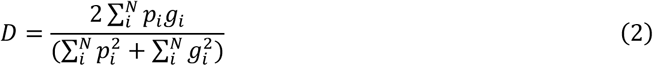

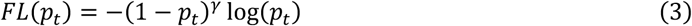

The model was trained for 120 epochs, taking over 18 hours on an Nvidia V-100 GPU. For learning rate scheduling, we followed previous works (*57*) and implemented learning rate warmup and cosine learning rate decay. During the first five epochs of training, the learning rate will increase linearly from 0 to 4×10^−3^. After that, we decreased the learning rate at each epoch following the cosine function. Based on our experiments, we ended up relaxing the learning rate decay such that the learning rate in the final epoch will be half of the initial learning rate instead of zero (*57*). We plotted the model’s loss value on both the training dataset and the validation dataset after each epoch (Fig. 2B) and picked the model checkpoint with the lowest validation loss as our final model to avoid overfitting. All the deep learning code was implemented using Python 3.8 and TensorFlow 2.3.

### Post-processing

We evaluated the performance of our trained E-U-Net on an unseen test dataset and reported the precision, recall, and F-1 score for each category: G1, S, G2/M, and background, respectively (Fig. S2). The pixel-wise confusion matrix indicated our model achieved high performance in segmenting the cell bodies from the background. However, since this pixel-wise evaluation overlooked the biologically relevant instance, i.e., the number of cells in each cell cycle stage, we performed an extra step of post-processing to evaluate that.

We first performed connected-component analysis on the raw model predictions. Within each connected component, we applied a simple voting strategy where the majority label will take over the entire cell. Figure S3A-B illustrate this process. We believe enforcing particle-wise consistency, in this case, is justified because it is impossible for a single cell to have two cell cycle stages at the same time and that our model is highly accurate in segmenting cell bodies, with over 0.96 precision and recall (Fig. S2). We then computed the precision, recall, and F-1 score for each category on the cellular-level. For each particle in the ground truth, we used its centroid (or the median coordinates if the centroid falls out of the cell body) to determine if the predicted label matches the ground truth. The cellular-wise metrics were reported in Fig. 4B.

Before using the post-processed prediction masks to compute the area and dry mass of each cell, we also performed hole-filling as we did for the ground truth masks to ensure the values are accurate (Fig. S3C).

## Acknowledgments

This work is sponsored in part by the National Science Foundation (0939511, 1353368), and the National Institutes of Health (R01CA238191, R01GM129709). C. H. is supported by the National Institutes of Health (T32EB019944). M.E.K. is supported by a fellowship from the Miniature Brain Machinery Program at UIUC (NSF, NRT-UtB, and 1735252).

## Data availability

All data required to reproduce the results can be obtained from the corresponding author upon a reasonable request.

## Code availability

All the code required to reproduce the results can be obtained from the corresponding author upon a reasonable request.

## Author Contributions

Y. H. and M. E. K. performed imaging. S. H., Y. H., and N. S. performed the deep learning development. Y. L. prepared cell cultures. Y. H. and C. H. performed analysis. Y. H., S. H., Y. L., C. H. and G. P. wrote the manuscript with inputs from all authors. M. A. A. supervised the deep learning. G. P. supervised the project.

## Supplementary Materials

**Fig. S1.**
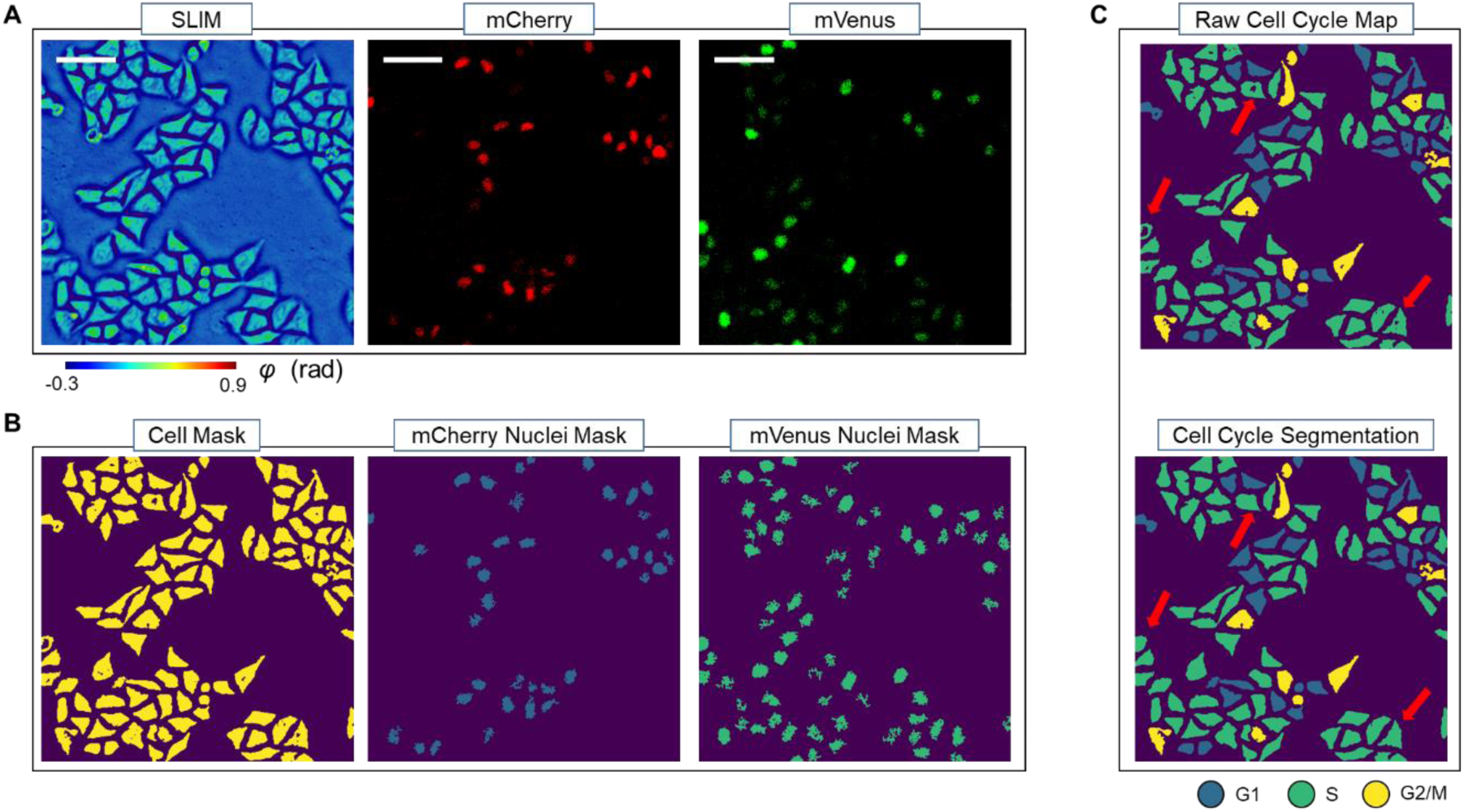
Ground truth mask generation workflow. (**A**) Images from the SLIM channel (left), mCherry channel (middle) and the mVenus channel (right). (**B**) Preliminary masks generated from the SLIM and fluorescence images using adaptive thresholding. (**C**) Combing three masks in b. Holes in cell masks were removed during analysis to avoid errors in cell dry mass and area. Scale bar is 100 μm.

**Fig. S2.**
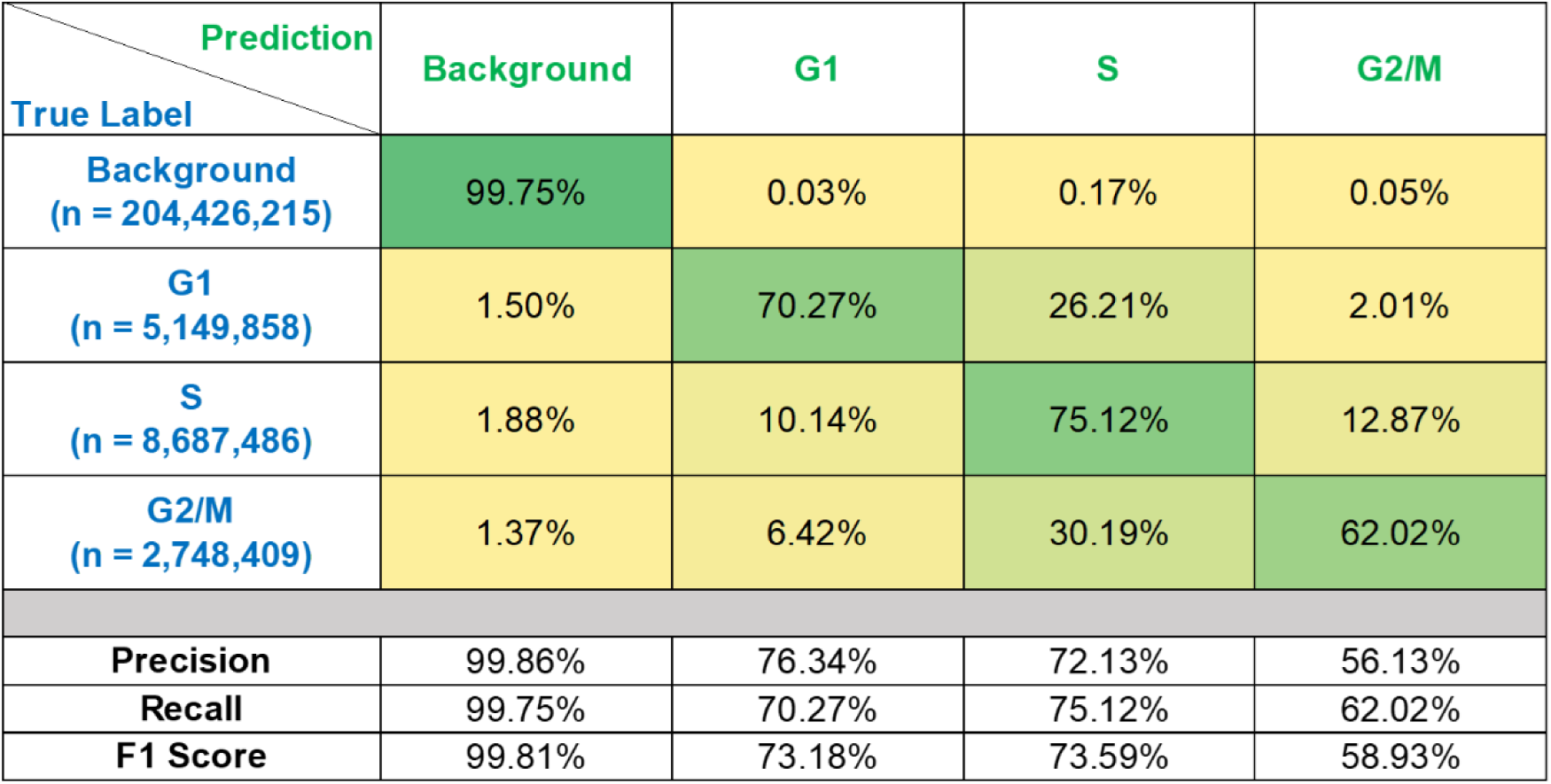
PICS performance evaluated at a pixel level.

**Fig. S3.**
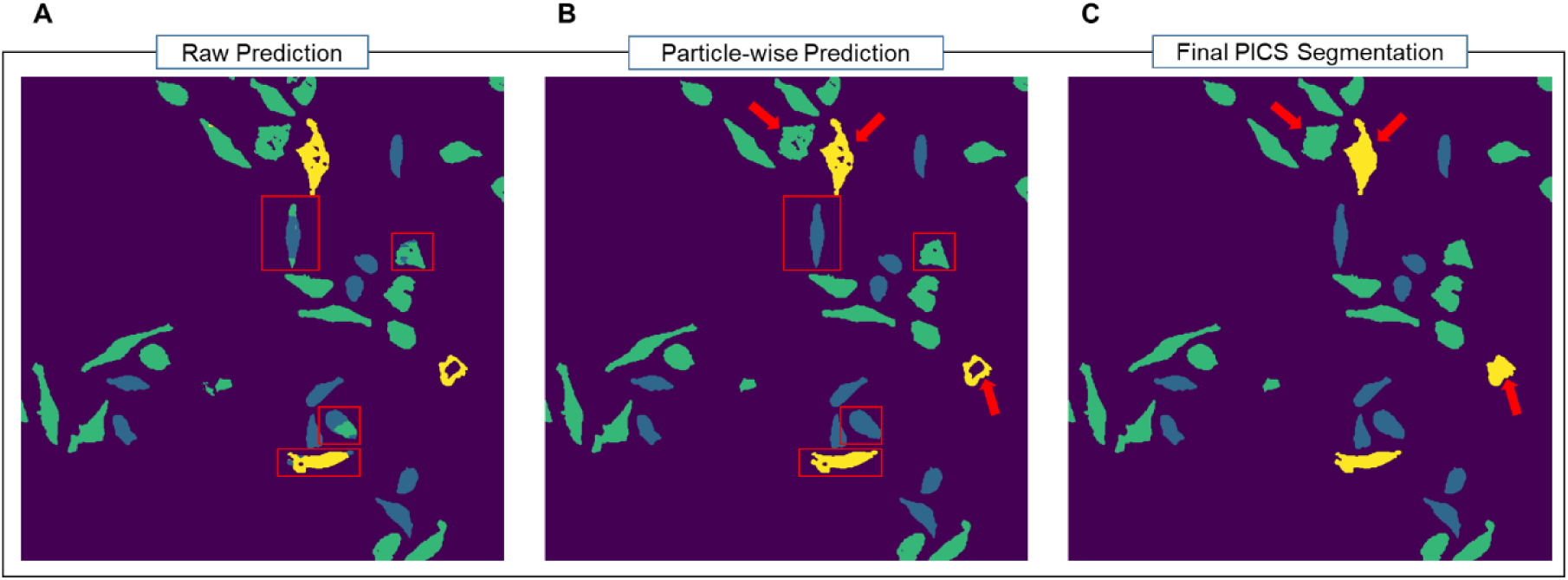
Post-processing workflow. (**A**) Raw prediction from PICS. (**B**) Prediction map after enforcing particle consistency and removing small particles. A few examples were shown in the red rectangles. (**C**) Prediction map after filling in the holes in the masks. Masks at this stage were used for analysis.

